# Implantable electrical stimulation bioreactor with liquid crystal polymer based electrodes for enhanced bone regeneration at mandibular large defects in rabbit

**DOI:** 10.1101/402719

**Authors:** Chaebin Kim, Hoon Joo Yang, Tae Hyung Cho, Beom Seok Lee, Tae Mok Gwon, Soowon Shin, In Sook Kim, Sung June Kim, Soon Jung Hwang

## Abstract

The osseous regeneration of large bone defects is still a major clinical challenge in maxillofacial and orthopedic surgery. Our previous studies demonstrated that electrical stimulation (ES) with biphasic current pulse showed proliferative effects on bone cells and enhanced secretion of bone-forming growth factors. This study presents an implantable electrical stimulation bioreactor with electrodes based on liquid crystal polymer (LCP), which has excellent bone-binding property. The bioreactor was implanted into a critical sized bone defect and subjected to ES for one week, where bone regeneration was evaluated four weeks after surgery using micro-CT. The effect of ES via bioreactor was compared with a sham control group and positive control group that received recombinant human bone morphogenetic protein (rhBMP)-2 (20 μg). New bone volume per tissue volume (BV/TV) in the ES and rhBMP-2 groups increased to 171% (*p* < 0.001) and 210% (p < 0.001), respectively, compared to that in the sham control group. In the histological evaluation, there was no inflammation within bone defects and adjacent to LCP in all groups. This study showed that the ES bioreactor with LCP electrodes could enhance bone regeneration at large bone defects, where LCP can act as a mechanically resistant outer box without inflammation.

## 1. Introduction

Regeneration of large bone defects has been a challenging issue in maxillofacial and orthopedic surgery. Extensively researched strategies include tissue engineering with cells, scaffolds, growth factors, mechanical environment, and biophysical stimulations [8,13,24]. Direct implantation of precultured stem cells is a straight-forward and effective method [27,58]. However, stem cell application for bone regeneration requires multiple complicated procedures such as tissue harvesting, separation of stem cells, cultultivation of cell for proliferation and differentiation, which are accompanied by long culture time, high cost and infection risk [6]. Moreover, the efficacy of stem cell transplantation is low because of low survival rate after transplantation [1]. Bone regeneration with osteoconductive cell-free substances such as allogenic, xenogenic or alloplastic bone grafts showed low osteogenesis efficacy with long regeneration time [20,43]. Bone regeneration with a growth factor such as recombinant human bone morphogenetic protein (rhBMP)-2 showed enhanced bone quantity with reduced regeneration time in the clinical application [16,38,56]. However, cell proliferation is not stimulated by rhBMP-2, while numerous cells are required for bone regeneration at large bone defects. Another relevant factor is biophysical stimulation under a suitable environment [19,35].

Bone morphogenetic protein (BMP) is a well-known osteoinductive signaling chemokine that causes bone formation by recruiting stem cells and differentiating them into bone-forming cells [28]. Among the BMP family, rhBMP-2 is widely used for bone tissue engineering because of its effectiveness in bone formation and its commercial accessibility. Despite its efficacy, rhBMP-2 is rapidly diffused when injected *in vivo*, hence, often high dose is required to obtain sufficient osteoinductive effect [53]. However, high dosage of rhBMP is associated with soft tissue hematoma swelling, radiculopathy, cyst-like bone formation, bone resorption, and inflammation [9,36,46].

To overcome the aforementioned limitations in bone regeneration, tissue prefabrication or prelamination techniques have been introduced for the regeneration of large bone defects. First, an avascular scaffold is vascularized after its implantation into well-vascularized tissue for a period; then, the vascularized scaffold with vessel pedicle is transferred to the tissue defects [18]. Based on this regenerative concept, tissue engineering components (scaffold, cells and growth factors) are implanted into an adequate anatomical site that can provide regenerative microenvironments with well-vascularized tissue, and the components are cultivated within living body tissue using intracorporeal self-regenerative function that acts as an *in vivo* bioreactor [18]. However, most large bone defect sites have reduced vascularity owing to the previous operations, infection or trauma, and thus they cannot provide optimal regenerative microenvironments. Therefore, new bone tissue should be first engineered at a well vascularized ectopic site, and, then, transferred to the bone defect site with vascular pedicles in the second surgery. Instead of such uncomfortable second surgery for surgeons and patients, it is preferable that regenerative microenvironments at bone defect sites can be improved by an appropriate physical stimulation such as electrical stimulation (ES) [33,34]; thus, tissue engineering components without vascular pedicles can be directly transferred into the osseous defect site.

ES modulates osteogenic activities of the stem cells, which can sense and react to the electric field [7]. ES has a positive effect on proliferation of bone cells, bone mineralization and expression of bone related growth factors and chemokine genes [3]. Recently, biphasic electric current stimulation was proposed to be used in bone regeneration for the safety of ES by achieving charge balance [29]. The biphasic electrical current stimulation on three dimensionally cultured mesenchymal stem cell accelerates the expression of chemokine receptors related to migration of the bone cells to the bone defect [34]. In our previous report, the feasibility of intracorporeally installed electrical stimulators as a form of *in vivo* bioreactor for enhanced bone regeneration at large mandibular bone defects was demonstrated [34]. However, the flexible electrodes did not have enough mechanical strength to maintain defect space against soft tissue tension after wound closure and external force during bone regeneration. Moreover, electrodes should be eliminated in the second surgery, because the electrodes used in the previous study were not integrated to the bone and may hinder bone union owing their area covering the cross-section of the defect.

Recently, liquid crystal polymer (LCP) has been used as a substrate for electrodes in neural prosthetic devices [22,52]. LCP is a biomaterial that has a mechanical strength similar to that of bone, an excellent affinity to bone and can be thermally deformed easily [25]. It can be effectively applied as an outer box or a scaffold for bone regeneration with ES in the orthopedic and maxillofacial field, because it can be integrated into regenerated bone, therefore its removal is not necessary. On this basis, we developed an implantable bioreactor with LCP electrodes, where LCP can act as a mechanically resistant outer box in bone defect. The aim of the present study was to evaluate the regeneration of new bone with or without ES, compared to that by rhBMP-2 without ES, at a large bone defect of the mandibular body in rabbits.

## 2. Methods

### 2.1 Fabrication of bioreactor

The bioreactor consisted of an LCP outer box containing gold electrodes, an electrical stimulator with a battery that produced biphasic electrical current pulses, and absorbable collagen sponges (ACSs) that were located within the LCP outer box. The LCP outer box was designed to fit the rabbit mandibular bone defect and its mechanical strength was sufficient to maintain the defect space necessary for bone regeneration. It was composed of three walls: the buccal, lingual and bottom walls; therefore, the residual three walls (upper, mesial and distal wall) in the hexahedral bone defect were covered with autogenous bone walls after mandibular bone resection, from which cell migration, angiogenesis and new bone ingrowth into the defect space could occur. Two patterned gold plates were embedded inside the LCP outer box to make the electric current flow across the bone defect (Fig 1A-D). ACSs were placed inside the LCP for osteoconductive bone regeneration (Fig 1E). The electrical stimulator, with a battery that was implanted in the submandibular area was connected with the gold electrodes of the LCP outer box through the lead that was fabricated together with the electrodes (Fig 1F).

**Fig 1.**
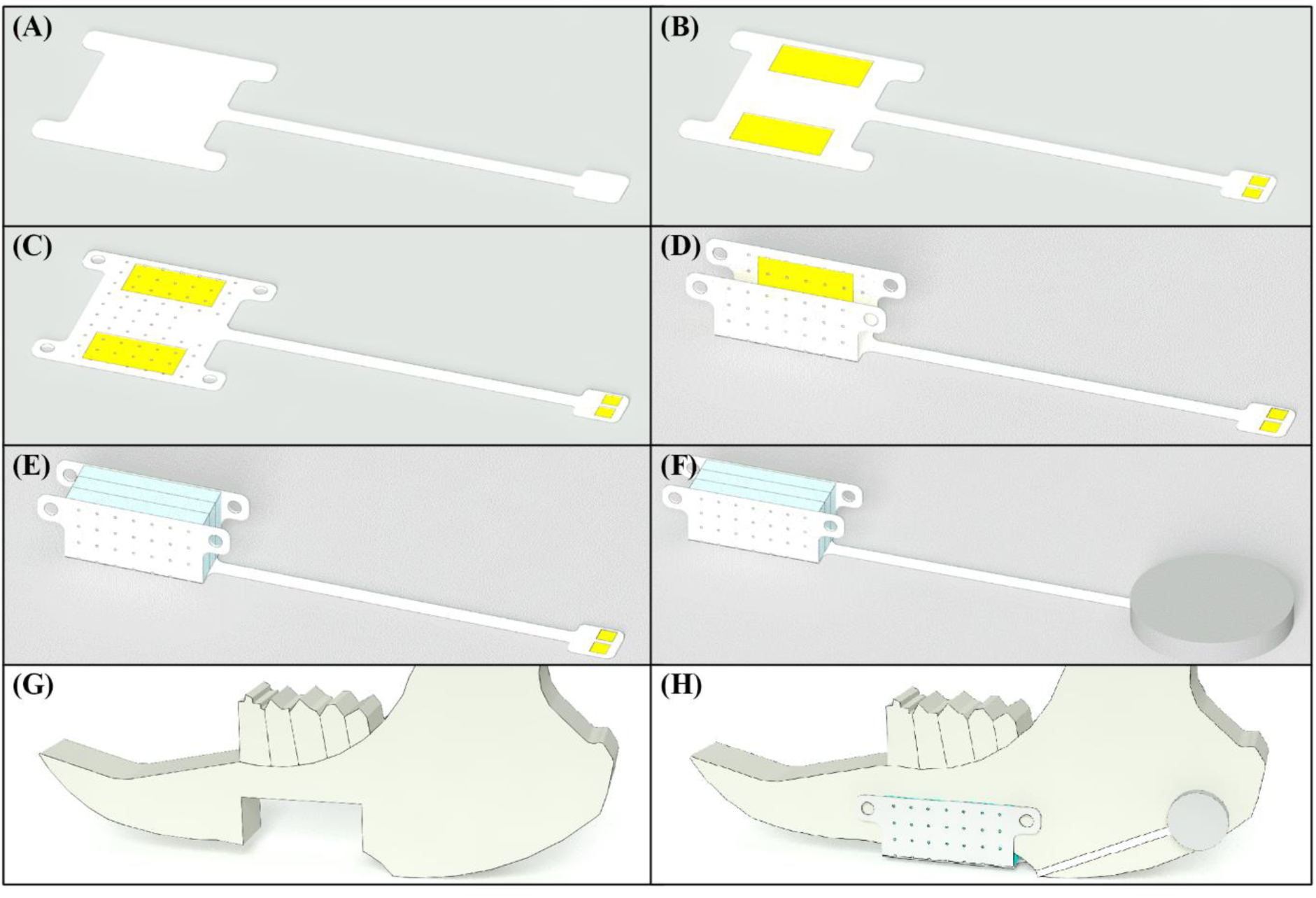
Fabrication of the bioreactor and process of the animal experiment: (a) LCP film is prepared as a substrate of the bioreactor, (b) gold electrodes are patterned on the substrate, (c) macro- and micro-holes are fabricated through the patterned film, (d) the film is deformed to be conformal to the bone defect, (e) collagen sponge is mounted in the space between the electrodes, (f) electric stimulator is connected to the extension plate, (g) mandibular bone defect is created by osteotomy, (h) the bioreactor is implanted to the bone defect and fixed with screws

#### 2.1.1 LCP outer box with gold electrodes

The LCP outer box was fabricated using LCP films (Vecstar, Kuraray, Tokyo, Japan) (Fig 2). The LCP outer box was made as follows. First, an LCP film was attached on a silicon wafer using a spin-coated silicone elastomer. Gold and titanium seed layer were deposited on the film using e-gun evaporator (ZZS550-2/D, Maestech, Pyeongtaek, Korea). The LCP film was dry etched with oxygen gas using plasma etcher (PlasmaPro System100 Cobra, Oxford Instruments, Abingdon, United Kingdom) for better metal adhesion. Photoresist (AZ4620, MicroChemicals, Ulm, Germany) was spin coated on the film for photolithography. The photoresist was patterned by photolithography using mask aligner (MA-6, SUSS MicroTec, Garching bei München, Germany). On the exposed gold layer, an additional gold layer with thickness of 5 μm was electroplated, and then photoresist and metal seed layers were wet-etched using photoresist remover, a mixture of nitric acid and hydrochloric acid, and hydrofluoric acid solution. Then, the LCP film was detached from the silicon wafer, and laminated with bare LCP films for insulation and thickness-control using heat press (model 4330, Carver, Wabash, USA). The outline of the film was cut with a laser, and the electrode site was ablated with a laser (355 nm UV, Samurai system, DPSS Lasers, Santa Clara, USA).

**Fig 2.**
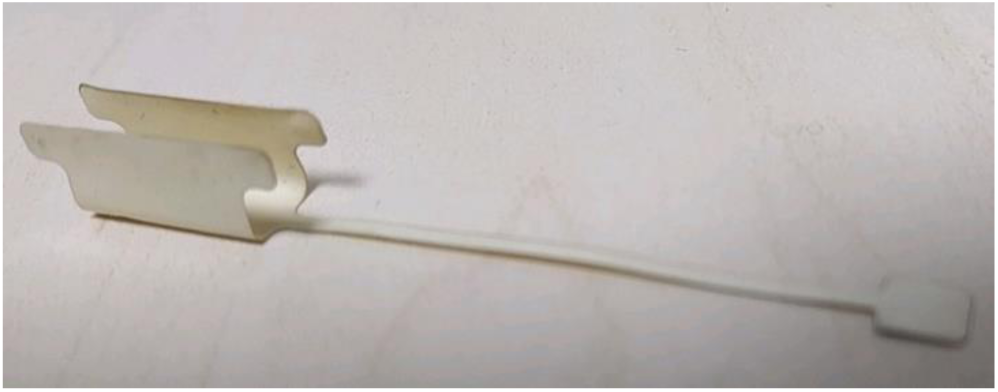
Fabricated LCP outer box whose shape and size are tailored to the target bone defect: the structure of the outer box is half cylindrical shell whose cross-section is U-shape. Size of the outer box is 15 mm in length, 7 mm in height, 6 mm in width. Thickness of the shell is 0.35 mm and interconnection line of 5 cm-length is extended for connection to electric stimulator

This LCP film was then thermoformed using prefabricated molds into a shape that could adapt to the rabbit mandibular bone defect. Two molds were fabricated: the outer mold was designed to possess the shape of the rabbit mandibular body with the created bone defect, and the inner mold was designed to deliver pressure to the LCP film. For the deformation, the LCP film was fixed in the outer mold and heated using the heat press. After the temperature reached 230°C, the inner mold was inserted in the outer mold to deform the LCP conformably to the surface of the outer mold. The temperature was maintained for 15 minutes before the LCP was cooled down. Then, the LCP was detached from the mold and treated with oxygen plasma followed by cleaning in ethyl alcohol in 60°C and isopropyl alcohol in 70°C.

After these procedures, the LCP electrodes possessed an identical U shape to the mandibular body surface shape prior to surgical resection, with dimension of 15 mm in length (anterior-posterior), 7 mm in height, and 6 mm in depth. The thickness of the LCP was adjusted to be 325 μm to provide sufficient mechanical strength for the maintenance of the defect space. On the inner buccal and lingual surface, gold electrodes were patterned to perform the ES at the bone defect space. The dimension of gold electrodes on both surfaces was 10 x 5 mm^2^. To facilitate ingrowth of new vessels and cells for transporting nutrition across the bioreactor, multiple micro-holes (500 μm in diameter) were made, with 2-mm distance between micro-holes. Four macro-holes were made at the upper corners of buccal and lingual LCP sides for the stabilization of the LCP with screws to mandibular bone. An extension plate with 2 mm width and 50 mm length was made for connection with the electrodes from the electric stimulator.

#### 2.1.2 Electric stimulator

Electric stimulator was designed using a custom designed chip (CMOS 0.18 µm foundry, Samsung electronics, Suwon, Korea) previously fabricated [34]. The circuit configuration of the stimulation chip was described in the previous study [33,34]. The integrated chip was wire bonded on the printed circuit board with additional components soldered [33]. The biphasic electric current pulses were generated by the electric stimulator, where the parameters of the pulses were set as amplitude of 20 μA, duration of 100 μs, and frequency of 100 Hz. The electric stimulator was connected with the LCP electrode and coin battery (CR1620, Panosonic, Kadoma, Japan) using wire and silver paste. The coin battery and the electric stimulator were packaged using silicone elastomer (MED-6215, Nusil, Carpinteria, USA) for the implantation into the submandibular area.

#### 2.1.3 Scaffold

Sterilized ACS sheets (Dalim Tissen Co. Ltd, Seoul, Korea) were cut into rectangle with dimensions of 15 mm width and 7 mm length. Three pieces of absorbable collagen sponge sheets were placed inside the LCP outer box for osteoconductive bone regeneration.

### 2.2 Surgical procedure

Fifteen male rabbits (New Zealandia rabbits, 3-3.5 kg weight) were used for the study. Animals were divided into three groups. The control group (n = 5) received neither ES nor rhBMP-2 after the surgery for the implantation of the bioreactor. In the ES group (n = 5), biphasic ES (10 ms time cycle, 100 μs duration, 20 μA amplitude) was applied to the bioreactor for one week immediate after the implantation of bioreactor. In the rhBMP-2 group (n = 5), rhBMP-2 (20 μg /defect) was injected into the defect space one week after the implantation of bioreactor without ES.

All animal experiments were performed in accordance with the “Recommendations for handling Regulation for Laboratory Animals for Biomedical Research” compiled by the Committee on the Safety and Ethical Handling Regulation for Laboratory Experiments in the School of Dentistry at Seoul National University. After general anesthesia using Ketamin (20 mg/kg, Yuhan, Seoul, Korea) and Xylazine (Rompun, 10 mg/kg, Bayer, Leverkusen, Germany) and skin disinfection with 10% Povidon-iodinie (Betadine, Purdue Pharma, Stamford, CN, USA), the mandibular body was exposed by incision into the skin and muscle dissection. Defect area to be resected (rectangular shape; 15 mm long, 7 mm high from the mandibular inferior border) was marked on the mandibular bone under the tooth and osteotomized by rotary drilling machine, where alveolar bone with teeth was preserved with continuity with the residual mandibular bone. Because mandibular stabilization with mini-plates after the resection of mandibular body including the alveolar bone with teeth was unstable [33], postoperative dislocation of anterior segment with loosening of mini-plates and screws occurred in our previous experiments due to the strong jaw movement. The bioreactors were positioned into the defect side, stabilized with screws on the neighboring mandibular bone (Fig 4), and the electric stimulator with battery coin were implanted at the submandibular area and then the incision sites were sutured layer by layer. The bioreactor and the electric stimulator remained at the bone defect for the duration of this experiment. Four weeks after the surgery, the animals were sacrificed and the defect sites, including neighboring bone, were resected for the micro-computer tomography (micro-CT) and histological evaluation. The resected bone tissues were fixed by dipping them in 10 % formalin solution.

**Fig 3.**
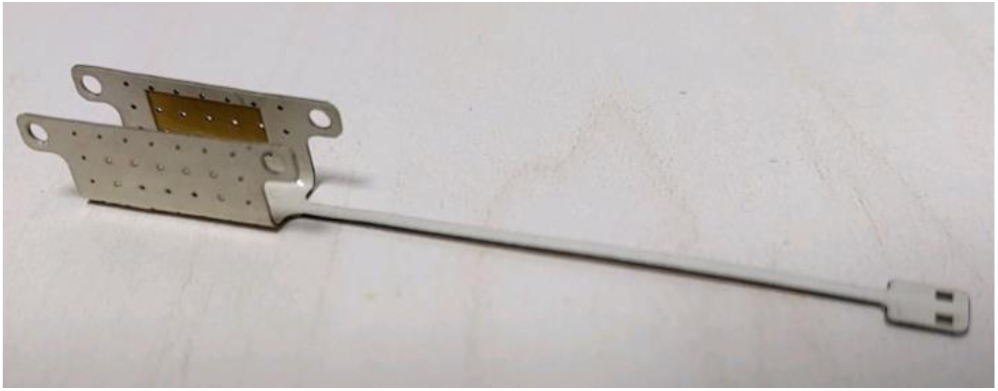
Fabricated LCP outer box with gold electrode and macro and micro holes: four macro holes are at the edge of the outer box to fix the device with screws at the bone defect. Micro holes are at the surfaces of the outer box to facilitate flow of body fluids between in and out of the box. Two gold electrodes are located at the side walls of the box for electric stimulation

**Fig 4.**
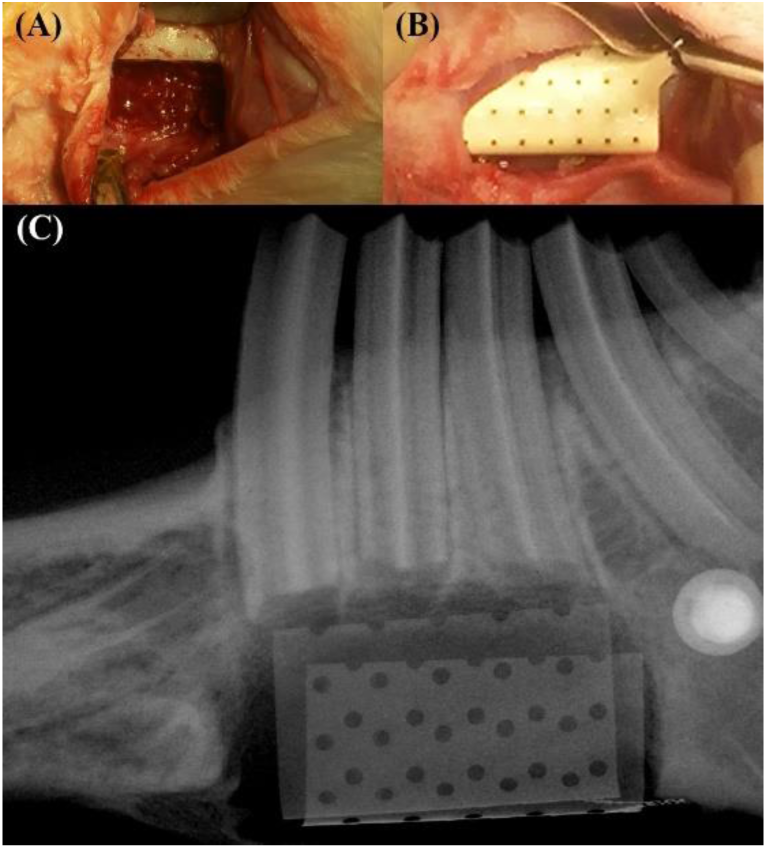
Intraoperative photographs and micro CT image after the implantation of bioreactor at the rabbit mandibular bone defect site: (a) bone defect was prepared by osteotomy at the mandibular body, (b) the bioreactor was implanted in the defect, where the LCP outer box is fixed on the neighboring bone with two screws, (c) micro CT image was taken to evaluate newly formed bone

### 2.3 Micro-CT

Micro-CT scan was taken for quantitative evaluation of new bone formation using Skyscan 1172 microfocus X-ray system (Bruker microCT, Kontich, Belgium). The SkyScan 1172 microfocus X-ray system is equipped with a microfocus X-ray tube with a focal spot of 2 mm, producing a cone beam that is detected by a 12-bit cooled X-ray camera CCD fiber optically coupled to a 0.5 mm scintillator. The resulting images were 2,000 x 1,048-pixel square images with an aluminum filter used to produce optimized images. A second-order polynomial correction algorithm was used to reduce the beam-hardening effect for all samples. Reconstructions and analyses were performed using NRecon 1.6.10.4 reconstruction (Bruker microCT, Kontich, Belgium) and CTAn 1.15.4.0 software (Bruker microCT, Kontich, Belgium), respectively. The reconstructed image was rotated three dimensionally for analysis using DataViewer (Bruker microCT, Kontich, Belgium). The 3D image was obtained using CTvol 2.3.1.0 (Bruker microCT, Kontich, Belgium).

In the reconstructed image, bone was isolated from soft tissue and noises by the adjustment of the brightness threshold of the pixel. The threshold between 115 and 230 was used to distinguish bone from noises, soft tissue, and electrode. The predefined defect area was selected as the region of interest (ROI) in two-dimensional images. The pixel zone representing ossification in the defined ROI was then reconstructed in 3D by setting the lower and upper ranges of the threshold using grayscale units. On each reconstructed BMP file, the bone volume per tissue volume (BV/TV), trabecular thickness (Tb.Th), trabecular separation (Tb.Sp), and trabecular number (Tb.N) were obtained using the CTAn 1.15.4.0 software according to the manufacturer’s instructions. To measure the bone mineral density (BMD), attenuation data for ROI or VOI were converted to Hounsfield units and expressed as a value of the BMD using a phantom (Bruker microCT, Kontich, Belgium). This phantom contained rods of calcium hydroxyapatite (CaHA) having a standard density corresponding to rabbit bone, which ranges from 1.26 to 1.65 g/cm3. BMD values were expressed in grams per cubic centimeter of CaHA in distilled water. A zero value for BMD corresponded to the density of distilled water alone (no additional CaHA), and a value greater than zero corresponded to non-aerated biologic tissue.

Newly formed bone was evaluated by two different methods. First, it was done in the whole bone defect space. In the second method, it was measured in five subparts; superior-buccal part, inferior-buccal part, superior-lingual part, inferior-lingual part, and central part, as shown in Fig 5. The upper boundaries of superior-buccal and superior-lingual parts were located at the upper surface of the bone defect, and the lower, buccal and lingual boundaries of each subparts were positioned at the lower, buccal and lingual surfaces of the LCP outer box. The central subpart was in the middle part in terms of the buccolingual width and the superior-inferior height of the whole bone defect. The region of interest was 15 mm in length (anterior-posterior), 2 mm in width and 2 mm in height.

**Fig 5.**
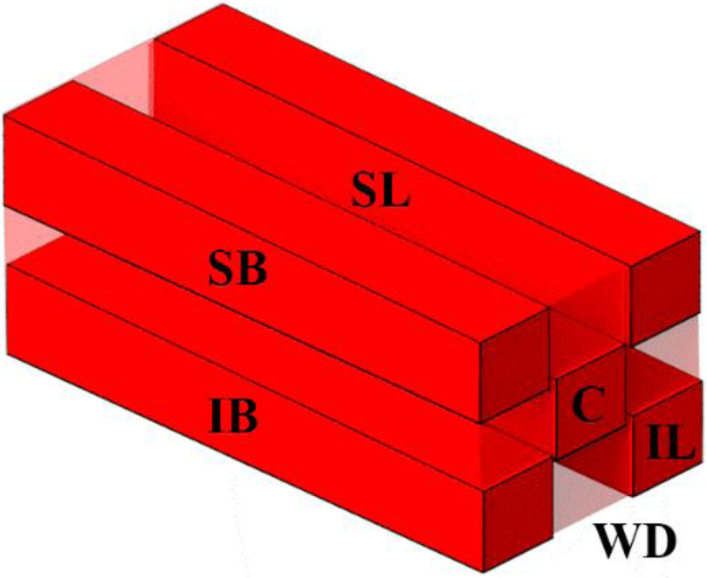
Region of interest (ROI) in the evaluation of new bone formation at the whole bone defect (WD) and at the five subparts: the size of ROI at subparts is 2 mm in width, 2 mm in height and 15 mm in anterior-posterior length. Subparts are superior-buccal (SB), inferior-buccal (IB), central (C), superior-lingual (SL), and inferior-lingual (IL) parts

### 2.4 Histological evaluation

After the CT image was taken, the resected bone tissues were further decalcified in an ethylenediaminetetraacetic acid solution (7 %, pH = 7.0) for 1-2 weeks. Then, the tissue samples were immersed in 70% ethanol and embedded in paraffin. For histochemical staining, decalcified paraffin sections were cleaned for 10 min with xylene and stained with Masson’s trichrome (MT). The decalcified specimens were sagittally cut at the middle plane of mandibular buccolingual width in the anteroposterior direction of the bone defect. Digital images of the stained sections were collected using a transmission and polarized light Axioscope microscope (Olympus BX51, Olympus Corporation, Tokyo, Japan).

### 2.5 Statistical analysis

The differences among bone parameters in the three groups were first tested using Kruskal Wallis H test and then analyzed using Mann Whitney U test. All statistical analysis was carried out using SPSS software (SPSS 25, IBM, NY, USA). A p-value < 0.05 was considered to indicate statistically significant differences.

## 3. Results

### 3.1 Macroscopic evaluation

Surgery sites including incision and bone defect area were healed without infection or wound dehiscence. LCP electrodes were maintained in the stabilized position without displacement. No inflammatory signs were seen between skin and bioreactor. All animals could be involved in the evaluation for micro-CT and histology.

### 3.2 Micro-CT analysis

#### 3.2.1 Bone regeneration in whole bone defect

The amount of newly formed bone volume per defect tissue volume (BV/TV) was 132 % greater in the ES group (11.9 ± 1.3 %) (p < 0.05) and 174% greater in the rhBMP-2 group (15.8 ± 1.5 %) (p < 0.01), compared to the control group (9.0 ± 2.1 %) (Fig 6, Table 1). The new bone was regenerated dominantly near the resected surfaces of the neighboring original mandibular bone. There was less new bone formation at the other areas, with especially poor bone regeneration at the central defect area (Fig 7). Concerning other parameters, Tb.Th and BMD were significantly higher only in the ES group (0.124 ± 0.007 mm and 1.581 ± 0.006 g/cm^3^, respectively) compared to those in the control group (0.113 ± 0.008 mm and 1.568 ± 0.010 g/cm^3^, respectively) (p < 0.05, both). Tb.N was significantly greater only in the rhBMP-2 group (1.34 ± 0.25 mm^-1^) compared to that in the control group (0.80 ± 0.15 mm^-1^) (p < 0.01). In the comparison between ES and rhBMP-2 groups, BV/TV and Tb.N were significantly higher in the rhBMP-2 group than in the ES group (p < 0.01 and p < 0.05, respectively), In summary, the rhBMP-2 group was superior in BV/TS and Tb.N compared to the ES group, while the ES group was superior to the rhBMP-2 group in terms of BMD and Tb.Th (Fig 6).

**Table 1.**
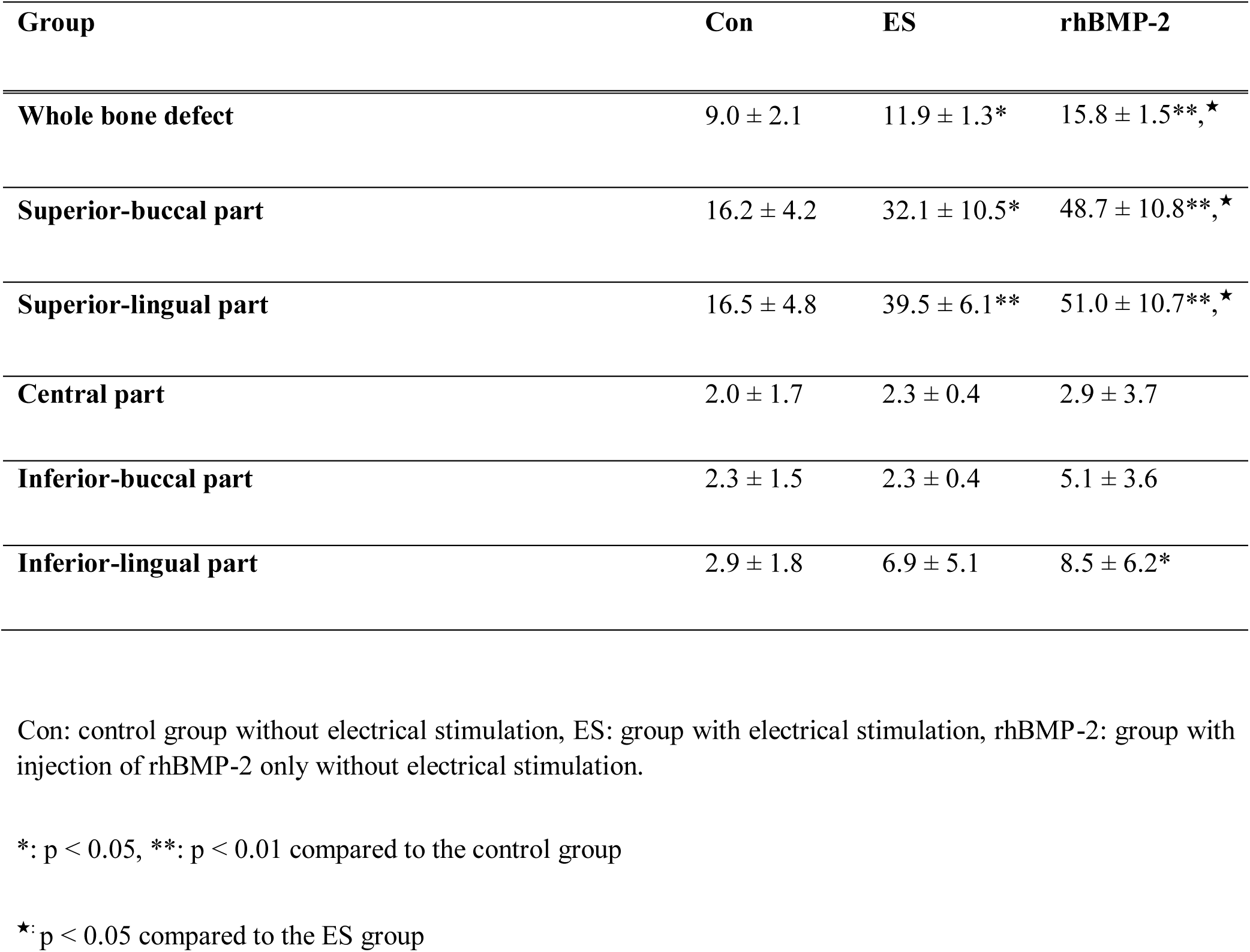
New bone volume per tissue volume (%) within the whole bone defect and five subparts.

**Fig 6.**
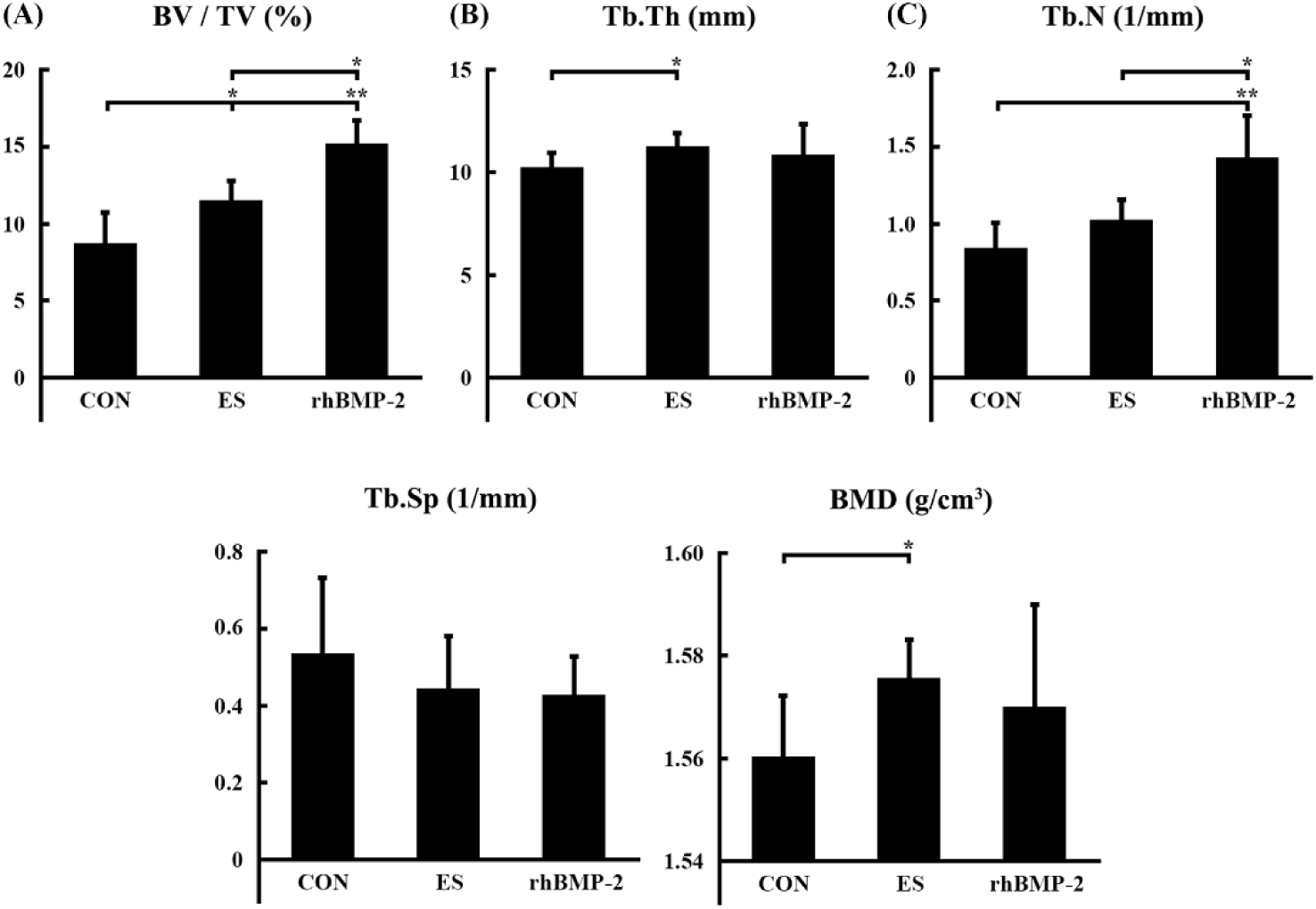
Analysis of newly formed bone within the whole bone defect in micro-CT evaluation: the analysis was performed in terms of (a) BV / TV, (b) Tb.Th, (c) Tb.N, (d) Tb.Sp, and (e) BMD. Con; control group without electrical stimulation, ES; group with electrical stimulation, rhBMP-2: group with injection of rhBMP-2 without electrical stimulation, BV/TV: bone volume/tissue volume, Tb.Th: trabecular thickness, Tb.N: trabecular number, Tb.Sp: trabecular separation, BMD: bone mineral density. Significant difference between the groups is indicated by *: p < 0.05, **: p<0.01

**Fig 7.**
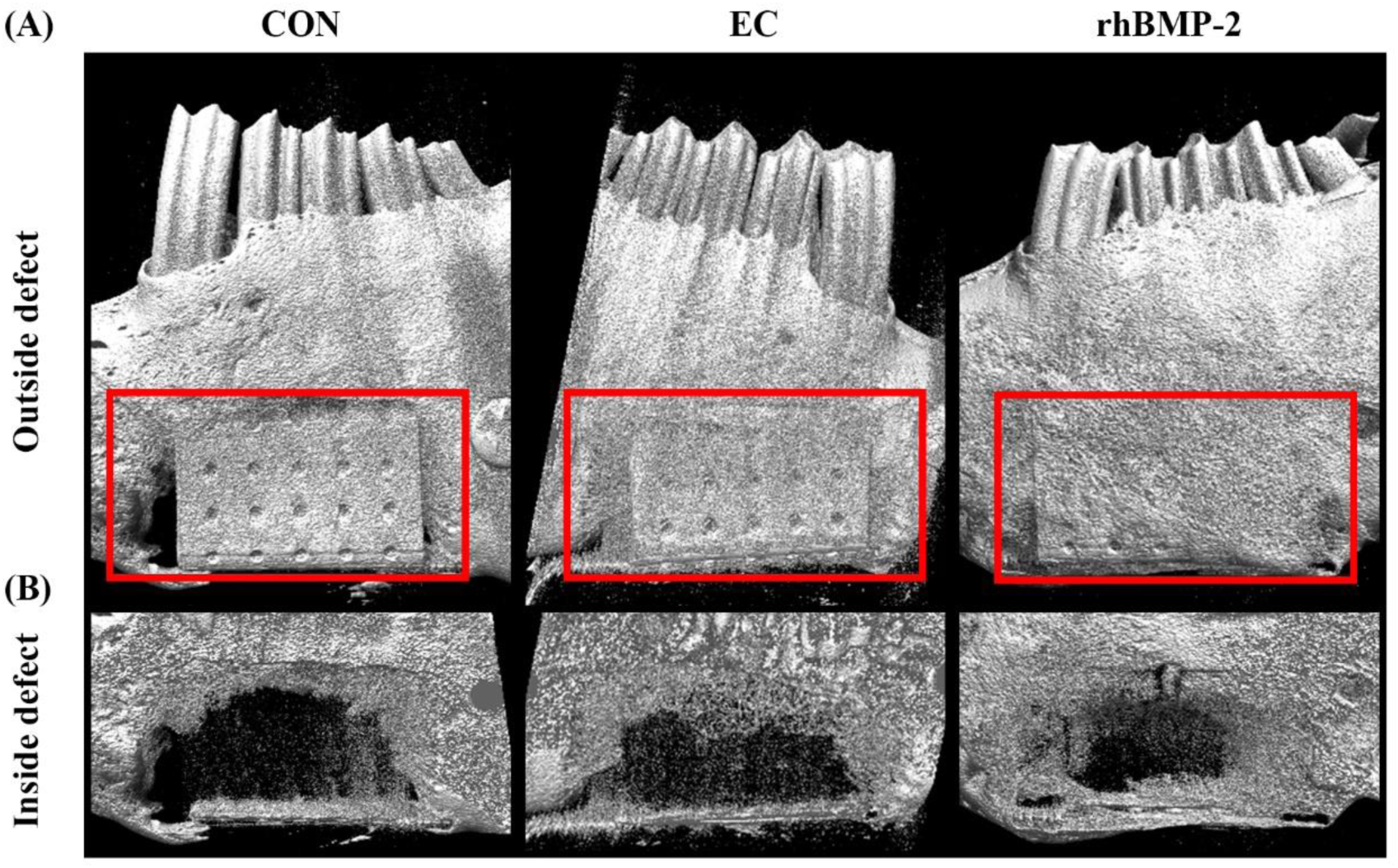
Micro-CT images of the mandibular bone after the experiment: micro-CT data was reconstructed in three-dimension to visualize the bone in the rabbit mandibular. The reconstructed image is manipulated by changing blending mode to show (a) mandibular external surface and (b) new bone within bone defect. Bone and gold electrode is depicted brightly, where the defect area is marked with red lines. Con; control group without electrical stimulation, ES; group with electrical stimulation, rhBMP-2: group with injection of rhBMP-2 only without electrical stimulation.

#### 3.2.2 Bone regeneration at five subparts

BV/TV was significantly higher at the superior subparts (34.1 ± 15.8 %) than at the inferior and central subparts (4.7 ± 3.5 % and 2.4 ± 2.2 %, respectively) (p < 0.01 for both), and significantly higher at inferior subparts than at central subparts (p<0.01). BV/TV at both buccal and lingual subparts (17.8 % and 20.9 %) was significantly greater than that at the central subparts (2.4 ± 2.2 %) (p < 0.01 for both), while there was no significant difference in new bone formation between the lingual and buccal subparts (Table 1)

The values of BV/TV (%) within five subparts are presented in Table. 1. In the superior subparts, ES and rhBMP-2 groups showed significantly greater BV/TV than in the control group. The superior-buccal subpart showed 205% increased BV/TV in the ES group (32.1 ± 10.5 %) compared to that in the control group (16.2 ± 4.2 %) (p < 0.05), while BV/TV in the rhBMP-2 group (48.7 ± 10.8 %) was 301% greater than that in the control group (p < 0.01). The superior-lingual subpart showed 239% increased BV/TV in the ES group (39.5%) compared to that in the control group (16.5%) (p < 0.01), while BV/TV in the rhBMP-2 group (48.7%) was 310% greater than that in the control group (p < 0.01) (Fig 8A, F).

**Fig 8.**
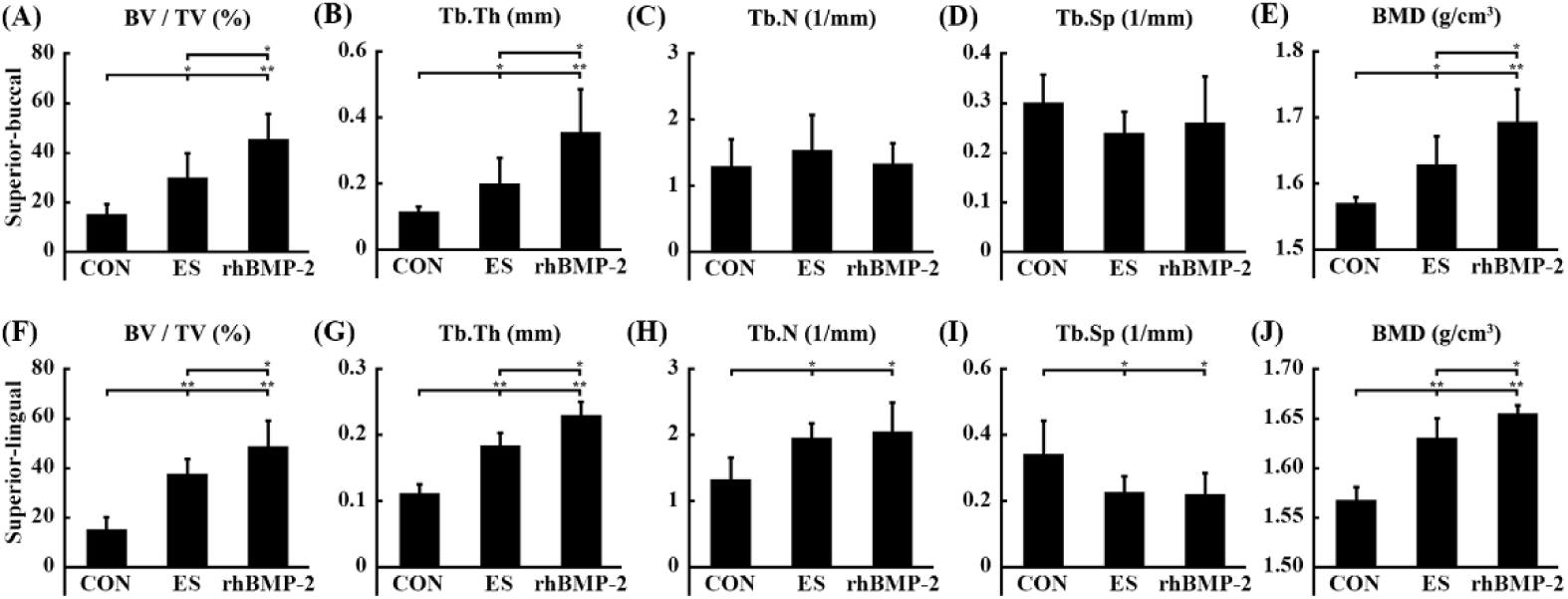
Micro-CT parameters of new bone at the superior-buccal and superior-lingual subparts in the three groups: the parameters at the superior-buccal subpart include (a) BV/TV, (b) Tb.Th, (c) Tb.N, (d) Tb.Sp, and (e) BMD. Also, the parameters at the superior-lingual subpart include (f) BV/TV, (g) Tb.Th, (h) Tb.N, (i) Tb.Sp, and (j) BMD. Con; control group without electrical stimulation, ES: group with electrical stimulation, rhBMP-2: group with injection of rhBMP-2 only without electrical stimulation. BV: bone volume, TV: tissue volume, Tb.Th: trabecular thickness, Tb.Sp: trabecular separation, Tb.N: trabecular number BMD: bone mineral density. Significant difference between the group is indicated by *: p < 0.05, **: p<0.01

In other parameters of the superior-buccal subpart, Tb.Th and BMD were significantly higher in the ES (0.21 ± 0.08 mm and 1.64 ± 0.04 g/cm^3^, respectively) and rhBMP-2 group (0.37 ± 0.13 mm and 1.70 ± 0.05 g/cm^3^, respectively) than that in the control group (0.12 ± 0.02 mm and 1.58 ± 0.01 g/cm^3^, respectively) (p < 0.05 between ES and control group, and p < 0.01 between rhBMP-2 and control group) (Fig 8B, E).

In the superior-lingual subpart, Tb.Th was significantly higher in the ES and rhBMP-2 groups (0.19 ± 0.02 mm and 0.24 ± 0.02 mm, respectively) compared to that in the control group (0.12 ± 0.01 mm) (p < 0.01 both). Tb.N was significantly higher in the ES and rhBMP-2 groups (2.05 ± 0.22 mm^-1^ and 2.14 ± 0.45 mm^-1^, respectively) compared to that in the control group (1.39 ± 0.34 mm^-1^) (p < 0.05 both). Tb.Sp was significantly lower in the ES and rhBMP-2 group (0.24 ± 0.05 mm and 0.23 ± 0.07 mm, respectively) compared to that in the control group (0.36 ± 0.10 mm) (p < 0.05 both). BMD was significantly higher in the ES and rhBMP-2 groups (1.64 ± 0.02 g/cm^3^ and 1.66 ± 0.01 g/cm^3^, respectively) compared to the control group (1.57 ± 0.01 g/cm^3^) (p < 0.01). (Fig 8G-J).

In the inferior and central subparts, there was no significant difference in BV/TV among three groups. The other values of micro-CT parameters and statistical significances are presented in Table 2.

**Table 2.**
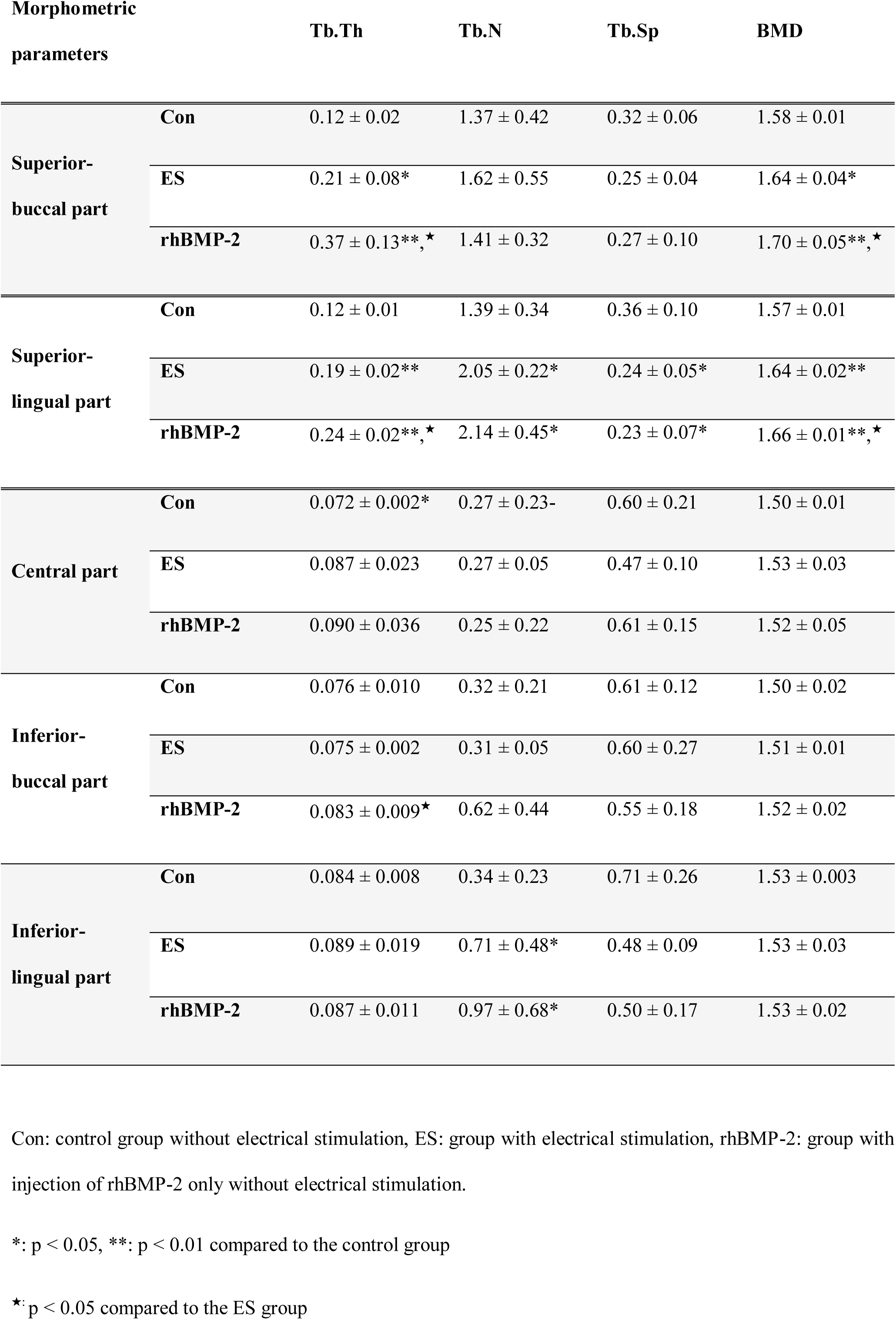
Trabacular bone thickness (mm), Trabacular bone number (1/mm), Trabacular bone separation (mm), and Bone mineral density (g/cm^3^) within the central and inferior subparts.

#### 3.2.3 Histological evaluation

The histological images stained in Masson’s trichrome showed the new bone regenerated in the bone defect area without inflammation in all groups, and none of the cases showed inflammatory cells at the interface between LCP and newly formed bone area (Fig 9, 10). Interestingly, thin new bone formation was observed along LCP at the inferior bone defect boundary (Fig 9 O, U and Fig 10 A, B, C), and especially thin osteoid layer was seen along LCP surface (Fig 9 B), which indicated a good biocompatibility of LCP for new bone formation. Excessive new bone formation could be seen at the inferior area of LCP in the rhBMP-2 group, probably due to the downward permeation of injected rhBMP-2 through the micro-holes of LCP (Fig 10C). In the control group, most new bone was thin and located near the neighboring bone surface defect with limited new bone ingrowth toward the central part, (Fig 9 A, D-H). In the ES and rhBMP-2 group, new bone was thicker, therefore, more ingrowth into the central part compared to that in the control group was observed (Fig 9 B, J-O, C, P-U).

**Fig 9.**
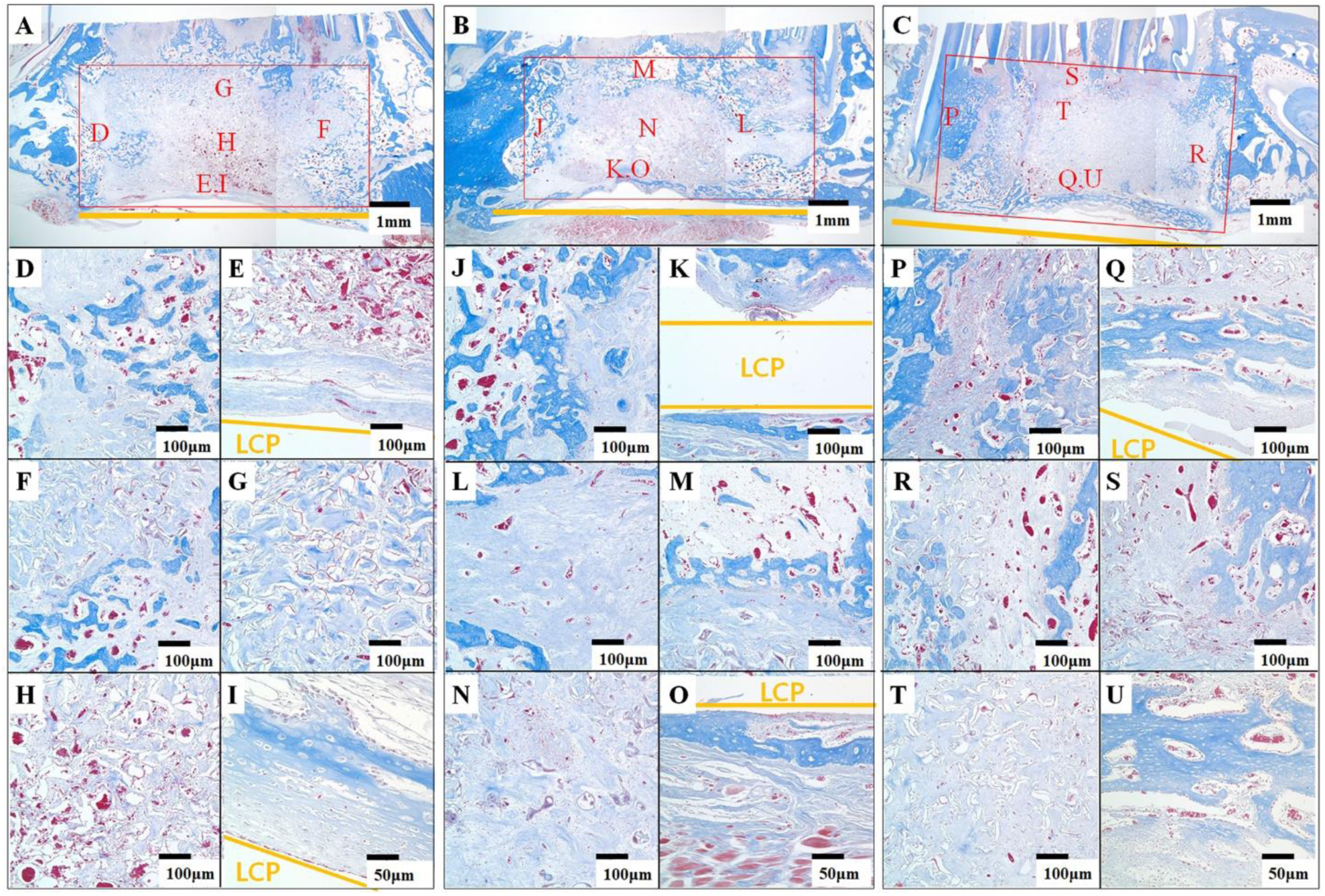
Histological images of the regenerated bone (Masson’s trichrome staining): the whole bone defect area is marked by a red rectangular line and the location of the bioreactor is marked in yellow in (a) (Control group), (b) (ES group), and (c) (rhBMP-2 group) (x 12.5). Each location of magnified images (D-H: control group, J-N: ES group, P-T: rhBMP-2, x 100) (I: control group, O: ES group, U: rhBMP-2, x 200) is depicted in A, B, C. The images are taken from different subparts of the defect such as (D, J, P) posterior, (E, I, K, O, Q, U) inferior, (F, L, R) anterior, (G, M, S) superior, and (H) central subpart. New bone has blue color, collagen has red color, extracellular matrix has light blue color, and red blood cells are red dots. Con; control group without electrical stimulation, ES; group with electrical stimulation, rhBMP-2: group with injection of rhBMP-2 without electrical stimulation

**Fig 10.**
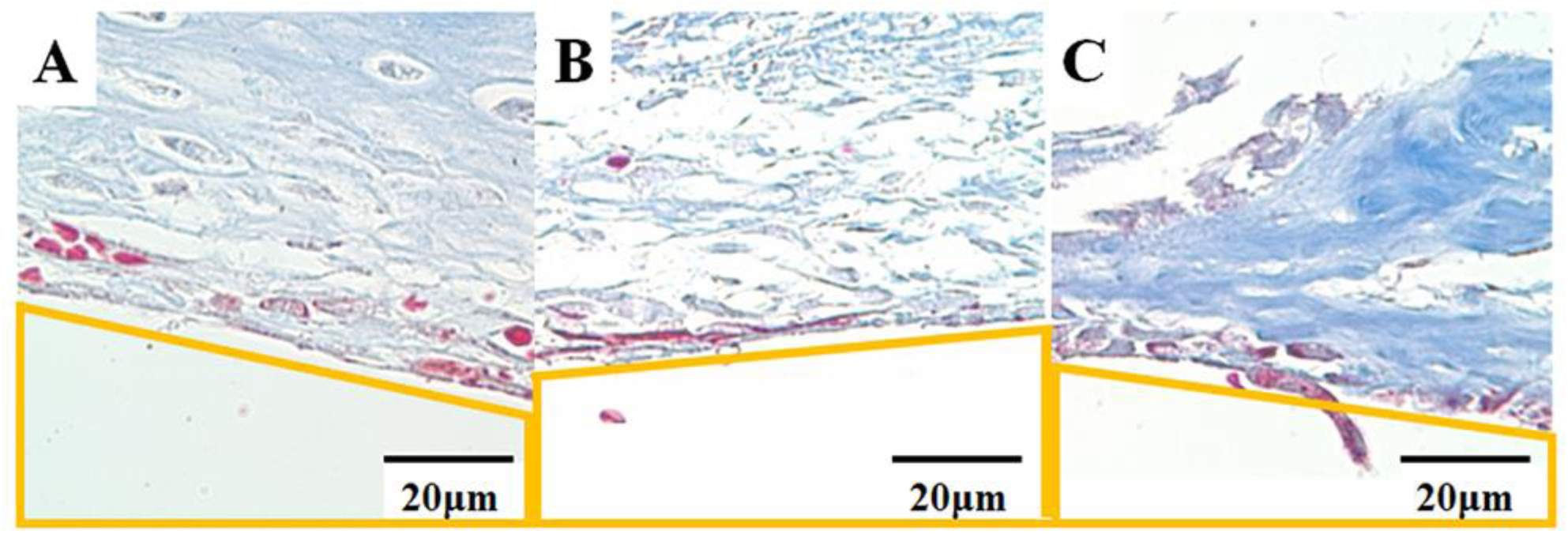
Histological images of the regenerated bone area contacted with the surface of the LCP outerbox in inferior-central region: location of LCP outer box is marked in yellow. There were no inflammatory reactions in newly regenerated bone in (a) CON, (b) EC, (c) rhBMP-2 group (Masson’s trichrome staining, magnification x500).

## 4. Discussion

This study addressed the effect of an *in vivo* bone bioreactor with ES on the bone regeneration at a large mandibular defect. We designed the bioreactor which was connected to custom-made electrodes based on LCP without stem cells or growth factor. LCP had multifunctional capacities and was biocompatible with bone, as well as mechanically stable to hold the shape of the bioreactor. Our bioreactor system is based on the previous reports that biphasic ES increased cell proliferation of human stem cells such as human mesenchymal stromal cell (hMSC) and several growth factors such as vascular endothelial growth factor (VEGF), BMP-2, and interleukin-8 (IL-8) [33,34]. These properties provide a great potential to create favorable microenvironments for bone regeneration. VEGF is a critical factor for angiogenesis, and IL-8 is a chemokine to recruit stem cells. This *in vivo* bioreactor system with ES benefits from one-staged direct application compared to a two-staged procedure where first the bioreactor is transplanted to the well-vascularized heterotopic sites prefabricated, and then, the regenerated tissue is transplanted to the orthotopic site [17,57].

The timing and duration of ES and BMP-2 treatment is an important aspect of experimental design for the optimal results. The effect of the ES in bone cells is different in different stages of cell maturation [10]. Further, the effect of the rhBMP-2 varies in different stages of bone cells [32]. The ES was conducted for one week after surgery, and BMP-2 was applied one week after surgery in this present study, based on reports that bone cell culture showed active proliferation but rare mineral deposition in the first week, while deposition was observed with declined proliferation after two weeks [11,37]. In addition, after fracture, cell proliferation lasted for 7 days, and then the cell differentiation and bone fusion occurred [55]. Furthermore, in the animal experiment with ES and BMP-2 [33], new bone formation was significantly higher in the group where BMP-2 application as delayed until one week after ES compared to that in the group with simultaneous application of ES and BMP-2. It seems that delayed BMP-2 application after the cell proliferation period is more efficient than immediate application after surgery.

Liquid crystal polymer (LCP) is a biocompatible material [2] and its mechanical strength is similar to that of the bone [23,25]. The tensile strength of LCP and cortical bone is 240 MPa and 150 MPa, respectively. The Young’s modulus of LCP and cortical bone is 2 GPa and 7 GPa, respectively. Because of the similarity in the mechanical strength and a good fixation-to-bone property [26], this material is considered adequate for orthopedic applications. Furthermore, it is chemically inert, allows hermetic sealing, and is compatible with MEMS microfabrication process with multilayer integration [15,21,40,51]. LCP-based packaging of a neural prosthetic device was studied [31,47,49]. Because of its thermo-forming property, it also allows the shape of the device to be formed to fit the target volume. This is the first report of an LCP-based electrode being used for bone regeneration. LCP based electrode was used for both ES and mechanical support of scaffolds. The electrode had good mechanical characteristic for bone regeneration, and the volume of newly formed bone was higher than with our previous ES device based on polyimide [33]. Further, histochemical staining showed no inflammation, suggesting that LCP is biocompatible. Micro CT images showed the bones formed on the outer and inner surfaces of the LCP. Histochemical staining demonstrated the new bone formation near the surfaces of the LCP. Our study suggested that electric stimulation with the implanted bioreactor based on LCP was effective in bone regeneration.

ES has been widely used as a method to enhance bone regeneration [39,42] and nerve regeneration [31,41]. ES increases proliferation, calcium deposition, and alkaline phosphate activity of bone related cells [12,30]. ES is known to open voltage gated calcium channels to increase intra-cellular calcium ions [5], which activates the cytoskeletal calmodulin [3]. Increased calcium level might lead to an increased nitric oxide levels, which acts in a physiological context through increased synthesis of cyclic guanosine monophosphate (cGMP) and subsequent activation of protein kinase G, which was associated with cell division and nucleic acid synthesis [48]. ES also made the cellular environment more suitable for osteogenesis [33]. The effect of ES included increased gene expression of type I collagen and prostaglandin [30], growth factors (BMP-2, IGF-1, VEGF), and chemokines with receptors [34]. ES activates mitogen-activated protein kinase (MAPK) and calcium channels, leading to increased induction of VEGF and BMP-2 of bone cells [29,30]. Additionally, ES increases transforming growth factor (TGF)-β1 through the calcium calmodulin pathway [60]. In the present study, micro-CT evaluation showed that the bone volume in the ES group was 132 % that of the control group. This result matched that of our previous experiment of ES in a dental implant, where the bone area in the ES group was 130 to 135 % of bone area in control group [29]. Even though ES could stimulate new bone regeneration, its effect on the amount of new bone formation was less than rhBMP-2 applied one week after surgery. It seems that enhanced cell differentiation by BMP-2 of small cell numbers is more efficient in bone regeneration than enhanced cell proliferation and differentiation by ES. However, Tb.Th and BMD were significantly higher only in the ES group compared to the control group, while there was no significant difference between the rhBMP-2 and control groups. These results suggest that new bone formation would be further enhanced in cases with initial ES for one week followed by rhBMP-2 application.

The effect of stimuli like ES or rhBMP-2 on bone regeneration can be different depending on the abundance of vascularity and stem cells. Corresponding to this expectation, new bone formation was predominant at the defect space adjacent to the existing host mandibular bone in the present study. Bone regeneration by ES was highest at the superior subpart of the bone defect than other subparts. This subpart was covered with more adjacent host bone surface including upper defect surface, providing more chance for stimulation of preexisting blood vessels and stem cells by proximal electrode. Based on the fact that electric current density is at is maximum near the electrode and gradually decreased as moving far from the electrode [50], and that the amount of the charge delivered is proportional to the amount of newly formed bone [4], more new bone could be observed near LCP. Considering that electric current can only flow through the conductive medium, it seems that host bone and newly regenerated tissue adjacent to bone defect are non-conductive substances. Blood clots and body fluid, which filled the bone defect space after surgery, may be electrically a conductive medium, but bone regeneration at the bone defect space far from existing host bone was poor, even with ES or rhBMP-2, because they could only be effective at the bone defect with infiltrated stem cells and ingrowing blood vessels. Vascularization is important for osteogenesis in large bone defect [14,45]. Biphasic electric pulse is effective for the increased secretion of VEGF from osteoblasts and stem cells[30], which is essential in angiogenesis [44] andstimulates new bone formation by osteoblasts [59]. Slow and steady release of VEGF promotes in bone formation [54], which can be effectively achieved through electric stimulation. On the other hand, the effect of ES or rhBMP-2 on bone regeneration can be influenced by the delivery efficacy of stimuli. According to the results of the present study and previous animal experimentation [33], initial ES followed by delayed BMP-2 application would be more efficient than ES or delayed BMP-2 application alone.

In conclusion, we verified the intracorporeally implanted ES bioreactor with LCP electrodes for bone regeneration, where LCP can act as a mechanically resistant outer box in bone defect without inflammation. The bioreactor system with ES significantly increased new bone formation compared to the control without ES at a large bone defect of rabbit mandible. The bioreactor system with ES was superior in BMD compared to rhBMP-2, while rhBMP-2 applied one week after surgery showed more enhanced osteogenesis than the bioreactor system with ES. Further experiments with other conditions of the bioreactor may be helpful to develop more effective bone regeneration.

## Acknowledgement

This work was supported in part by the Dentistry - Engineering Interdisciplinary Research Grant jointly funded by the School of Dentistry and College of Engineering, Seoul National University; and in part by a grant to CABMC project funded by the Defense Acquisition Program Administration (UD140069ID)

